# Astrocyte-Selective Wireless Deep Brain Stimulation by Magnetite Nanodiscs at Low Alternating Magnetic Field

**DOI:** 10.64898/2026.08.13.744740

**Authors:** Yi-Ting Cai, Jiao-Cheng Wang, Po-Han Chiang

## Abstract

Astrocytes are essential regulators of synaptic transmission, brain homeostasis and behavior, and direct astrocyte activation has emerged as a therapeutic target for a broad range of neurological disorders. However, selective activation of astrocytes in vivo currently requires viral transgene delivery, tethered fiber implants, or MRI-scale magnetic fields, limiting deployment across laboratories and translational settings. Here we introduce anti-GLAST-conjugated magnetite vortex nanodiscs (αGLAST-MND) that, together with a low-amplitude alternating magnetic field, enable wireless, transgene-free, cell-type-specific astrocyte stimulation. The torque generated by MND is sufficient to trigger mechanosensors without the strong spatial gradient required by prior magnetomechanical glial stimulation approaches. αGLAST-MND injected into the mouse dentate gyrus co-localized with the astrocyte membrane marker GLAST in vivo. Fiber-photometry in paired astrocyte (gfaABC1D-GCaMP6f) and neuron (Thy1-GCaMP6s) reporter cohorts showed that stimulation at 25 to 28 mT and 10 Hz evoked astrocyte Ca²⁺ responses 2-to 3-fold larger than non-magnetic hematite controls, while the neuron cohort showed no material-specific response. A single intracranial injection sustained stable astrocyte responses across at least five weeks and extended to seven weeks in a smaller follow-up cohort. Immunohistochemistry using the contralateral hemisphere as an internal reference confirmed that αGLAST-MND stimulation did not induce local c-Fos, consistent with the absence of downstream neuronal excitation. This approach uses a scalable and affordable magnetic system compatible with freely-behaving animals, opening opportunities for astrocyte-focused neuroscience research and for the future development of astrocyte-targeted therapeutics.

## 1. Introduction

Astrocytes are the most abundant glial cell type in the mammalian brain and are essential regulators of synaptic transmission, brain homeostasis and behavior. Beyond canonical functions such as glutamate uptake, extracellular potassium buffering, and maintenance of the blood-brain barrier, astrocytes actively shape neuronal circuit output through Ca²⁺-driven gliotransmitter release and metabolic coupling to neurons. Cell-type-specific studies in rodents have identified astrocyte contributions to sleep-wake regulation [1], learning and memory [2], sensory cortical processing [3], and striatal goal-directed behavior [4]. Aquaporin-4-mediated glymphatic clearance depends on astrocyte function to remove amyloid-β and other metabolic waste from the brain [5, 6]. On the disease side, dysfunctional astrocyte-to-neuron coupling drives chronic pain through spinal and supraspinal mechanisms [7], underlies major depressive disorder [8], contributes to epilepsy [9], and accelerates Alzheimer’s and Parkinson’s disease pathology [10]. Because astrocyte dysfunction spans neurological and psychiatric disease categories, astrocyte-targeted therapeutic interventions have received growing attention across CNS disorders [11, 12].

Direct astrocyte activation in vivo has been demonstrated with chemogenetic and optogenetic tools. Chemogenetic activation of striatal astrocytes partially rescues motor deficits in the parkinsonian mouse model [13]. Chemogenetic activation of hippocampal CA1 astrocytes enhances memory allocation [2]. Optogenetic activation of hypothalamic astrocytes increases sleep [1]. Optogenetic activation of cortical astrocytes modulates the response selectivity of visual cortex neurons [3]. These preclinical studies support astrocyte stimulation as a therapeutic modality complementary to neuron-targeted brain stimulation. However, chemogenetic approaches require viral transduction of a DREADD receptor and administration of an exogenous ligand, and optogenetic approaches require viral transduction of an opsin and a chronically implanted optical fiber for photostimulation. Both constraints limit clinical translation and restrict experimental flexibility. A wireless, transgene-free stimulation platform that engages astrocytes selectively *in vivo* would open a route to astrocyte-targeted neuromodulation in awake, freely-behaving animals and to eventual clinical deployment.

Current clinical transgene-free neuromodulation technologies, including deep brain stimulation (DBS), transcranial magnetic stimulation (TMS), electroconvulsive therapy (ECT) and transcranial focused ultrasound (FUS), have transformed the treatment of Parkinson’s disease, essential tremor, dystonia, treatment-resistant depression and obsessive-compulsive disorder [14, 15]. Each of these modalities delivers therapeutic benefit by perturbing local electrical activity within a defined brain region, and several are now standard of care for medication-refractory neurological disease. However, none can selectively engage a defined cell type. DBS electrodes activate every neuron, axon and glial process within the electric field; TMS depolarises cortical neurons and the glia surrounding them indiscriminately; ECT seizures spread through all cellular compartments; and FUS modulates whichever mechanosensors lie within its focal spot regardless of cell identity. As a consequence, the astrocyte-targeted therapeutic strategies described above cannot be selectively engaged by any clinically approved neuromodulation technology. Closing this gap therefore requires a different stimulation principle in which cell-type recognition occurs at the molecular interface rather than at the level of the electric field.

Wireless magnetic-nanoparticle neuromodulation directly addresses these constraints. Alternative magnetic fields (AMF) less than 1 MHz traverse biological tissue with negligible attenuation and require no implanted hardware [16, 17]. Magnetothermal [16, 18, 19] and magnetomechanical [20, 21, 22] modalities have been demonstrated in neurons, and magnetite vortex nanodiscs (MNDs) drive low-amplitude (< 50 mT) magnetomechanical activation of endogenous mechanosensitive channels in peripheral and central neurons [20, 22, 23]. All reported MND applications to date have targeted neurons, however, and no MND platform has combined this low-field regime with transgene-free, cell-type-specific targeting. The single prior magnetomechanical study on astrocytes used anti-GLAST antibody-functionalized iron oxide spheres actuated by either the fringe field of a 9.4 T MRI scanner or a purpose-built permanent magnet array [24]. The MRI-based approach restricted to institutions with dedicated imaging infrastructure. The permanent magnet array has a fixed geometry that is not scalable for freely-behaving recording chambers. In contrast, a recently reported open-source coil system delivers low-amplitude AMFs at bench-top scale for a total hardware cost less than 3,000 USD, and can be reconfigured for both freely-behaving animal chambers and in vitro study [25]. Astrocytes natively express a rich complement of mechanosensitive channels amenable to low-amplitude actuation, including TRPV4 [26], Piezo1 [27, 28], and TRPC1 [29]. Whether transgene-free, cell-type-specific astrocyte stimulation can be delivered by such affordable, scalable open-source hardware remains untested.

To wirelessly modulate astrocyte activity by magnetomechanical stimulation in a cell-type-specific manner, we conjugated magnetite vortex nanodiscs (MNDs) with an astrocyte-specific antibody, anti-GLAST, that binds the astrocyte plasma membrane but not neurons (Figure 1A). Under AMF, the MND-generated torque on the astrocyte membrane activates mechanosensitive ion channels. αGLAST-MNDs were injected into the dentate gyrus and stimulated through the open-source coil system described above [25]. Paired gfaABC1D-GCaMP6f (astrocyte reporter) and Thy1-GCaMP6s (neuron reporter) mice with chronic fiber-photometry recording showed that αGLAST-MND stimulation evoked reproducible astrocyte Ca²⁺ responses at 25 to 50 mT, produced significantly larger astrocyte than neuron Ca²⁺ responses, and maintained efficacy for at least 5 weeks from a single intracranial injection. The platform provides a wireless, transgene-free tool for astrocyte-targeted research on bench-top magnetic hardware, with potential translational applications to disorders in which astrocyte dysfunction is causally implicated.

**Figure 1.**
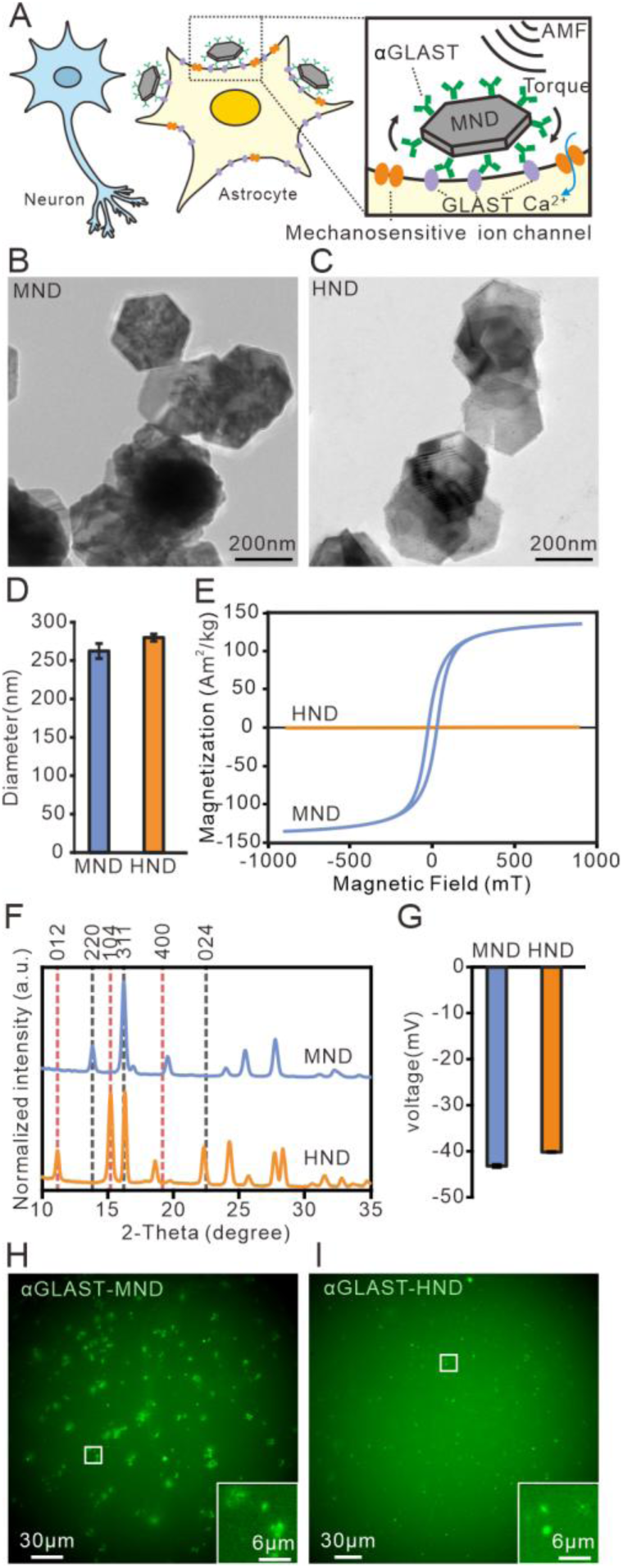
Characterization of astrocyte-targeting magnetic vortex nanodiscs. (**A**) Schematic of the platform: MND functionalised with an anti-GLAST antibody, specifically tethered to the astrocyte plasma membrane by binding to the GLAST on membrane. Inset, during AMF stimulation, the torque generated by MND triggers opening of mechanosensitive ion channel and cause Ca2+ influx. (**B**) TEM image of MNDs. (**C**) TEM image of HNDs. (**D**) Diameters of MND and HND quantified from TEM images (n = 6 for each). (**E**) VSM hysteresis loops for MND and HND. (**F**) Powder XRD patterns confirming full phase conversion of the α-hematite precursor (HND) to the magnetite (MND). (**G**) Zeta-potential of anti-GLAST-conjugated MND and HND particles in ddH2O (n = 3 for each). (**H**) Fluorescence image of αGLAST-MNDs with an Alexa Fluor 488-conjugated secondary antibody. (**I**) Same as (H) for αGLAST-HNDs.

## 2. Materials and Methods

### 2.1 Nanodiscs and antibody conjugation

Hematite (HND) and magnetite (Fe₃O₄; MND) vortex nanodiscs were synthesized, PMAO-coated, and conjugated with anti-GLAST (Miltenyi Biotec #130-095-822) or anti-Thy1 (clone G7) monoclonal antibody as previously described [23, 25] (Supplementary Methods S1-S2). HNDs served as the magnetic-effect control. Nanodiscs were characterized by TEM, VSM, XRD, DLS and ICP-OES (Supplementary Methods S3).

### 2.2 Animals

Adult male C57BL/6J (BioLASCO Taiwan) and Thy1-GCaMP6s (Jackson Laboratory, #024275) mice were used. All procedures were approved by the National Yang Ming Chiao Tung University IACUC (protocol #114009) and reported in accordance with ARRIVE 2.0. Mice were randomly assigned to MND or HND groups (Supplementary Methods S4).

### 2.3 Stereotactic injection and fiber implantation

Under isoflurane anesthesia, mice received a unilateral dentate gyrus (DG) injection of AAV5-gfaABC1D-cyto-GCaMP6f (0.6 μL, AP −1.9 / ML +1.2 / DV −1.9 mm relative to bregma). After ≥ 3 weeks of viral expression, αGLAST-MND or αGLAST-HND particles (2 mg mL⁻¹, 2 μL) were injected at the same coordinates and an optical fiber (200 μm core, 3 mm length, 0.48 NA) was implanted 100 μm dorsal to the injection site (Supplementary Methods S5).

### 2.4 AMF stimulation apparatus

Alternating magnetic fields (AMFs) were generated by an open-source air-core coil system as previously described [25]. Peak field amplitude was calibrated with a Hall-effect gaussmeter before every experimental cohort (Supplementary Methods S6).

### 2.5 Fiber-photometry recording and signal analysis

Fiber-photometry was performed in freely-moving awake mice within the stimulation chamber, with 410 / 470 nm alternating excitation at 15 fps per channel. Recordings were performed once or twice per week for 7 weeks post-injection. Traces were processed in Python; ΔF/F₀ = (F − F₀) / F₀ with F₀ defined as the mean of the lowest 5% of samples in the preceding 100 s. The primary metric was max change ΔF/F₀ during the 30 s stimulation window minus the mean 20 s pre-stimulus baseline. Details in Supplementary Methods S7-S8.

### 2.6 c-Fos experiment

Three weeks after unilateral DG injection of αGLAST-MND, αGLAST-HND, αThy1-MND or αThy1-HND (n = 6/group; contralateral hemisphere as internal reference), mice were stimulated at 28 mT × 10 Hz (10 cycles of 30 s on / 30 s off), perfused 90 min later, and processed for c-Fos / DAPI immunolabeling (Supplementary Methods S9-S10).

### 2.7 Statistical analysis

Sample sizes are reported in the figure legends and were based on effect sizes from analogous magnetomechanical studies [20, 22, 23, 24]. Per-mouse values were computed by averaging trials within each mouse × week × field combination.

Paired within-material baseline versus stimulation used the Wilcoxon signed-rank test; between-material comparisons of max change ΔF/F₀ used the Mann-Whitney U test; cell-type specificity was tested by two-way ANOVA (Material × Cell type). Effect size was quantified as Cohen’s d with two-sided 95% percentile bootstrap confidence intervals (10,000 iterations, mice resampled with replacement) [30]. All tests were two-tailed with α = 0.05. Analyses used Python 3.9 (scipy, statsmodels, DABEST). Data are mean ± SEM.

## 3. Results

### 3.1 Engineering of anti-GLAST-functionalized MNDs

MNDs and geometry-matched non-magnetic α-hematite disc controls (HNDs) were synthesised as previously described [22, 31]. Briefly, HND precursors were prepared by autoclave-assisted hydrothermal growth and MNDs obtained by subsequent reduction of hematite to magnetite (Fe₃O₄); HNDs (the un-reduced intermediate) served as the magnetic-effect control throughout all in vivo experiments. TEM confirmed the disc-like morphology of both particles (Figure 1B, C), with diameters of 262 ± 10 nm (MNDs) and 280 ± 5 nm (HNDs) (Figure 1D). VSM revealed the low-coercive-field vortex hysteresis of MNDs (saturation magnetisation Ms ≈ 135 Am² kg⁻¹) and the negligible antiferromagnetic response of HNDs (Ms ≈ 0.4 Am² kg⁻¹; Figure 1E). XRD verified full phase conversion of the α-hematite precursor to magnetite (Figure 1F). To confer cell-type-specific targeting, MNDs were coated with poly(maleic anhydride-alt-1-octadecene) (PMAO) and conjugated to a monoclonal anti-GLAST antibody by carbodiimide coupling. Zeta-potential measurements yielded negative surface charge for both αGLAST-MND (−43.2 ± 0.3 mV) and αGLAST-HND (−40.2 ± 0.2 mV) preparations (Figure 1G), consistent with successful covalent antibody attachment. An Alexa 488-conjugated secondary antibody labelled both αGLAST-MND and αGLAST-HND (Figure 1H, I), confirming successful antibody conjugation. Together these characterizations establish an antibody-functionalized vortex MND preparation, and a geometry-matched non-magnetic HND control, suitable for selective tethering to the astrocyte plasma membrane in vivo.

### 3.2 áGLAST-MNDs bind astrocyte-restricted GLAST epitopes *in vivo*

Because the anti-GLAST antibody targets the astrocyte plasma-membrane transporter GLAST/ EAAT1 [32], it cannot co-localize with the canonical astrocyte marker GFAP, which is an intracellular intermediate-filament cytoskeletal protein confined to the astrocyte soma and thick primary processes [33]. Astrocyte-restricted binding was therefore validated against a second astrocyte plasma-membrane glutamate transporter, EAAT2, which shares GLAST’s compartment and is expected to yield peak-aligned line-scan profiles. Cell-type specificity against neurons was independently assessed using the neuronal marker MAP2, a cytoplasmic microtubule-associated protein enriched in the neuronal soma and dendrites [34], expected to yield peak-alternated line-scan profiles against astrocyte-restricted GLAST.

Direct anti-GLAST immunostaining of dentate gyrus (DG) sections produced the reticulated, process-rich signal characteristic of astrocyte perisynaptic membranes and closely tracked EAAT2 across the molecular and granule cell layers, with line-scan profiles showing peak-aligned co-variation (Figure 2A, B, S1). The same anti-GLAST signal, co-stained with MAP2, showed peak-alternation, with MAP2-positive somatodendritic profiles occupying the intervening valleys (Figure 2C, D, S1). αGLAST-MNDs were injected into the DG and detected with a 2nd antibody targeting the MND-borne monoclonal (Figure 2E, G, S1). Line-scan profiles reproduced the direct-IHC pattern: peak-aligned co-variation with EAAT2 (Figure 2E, F, S1) and peak-alternation with MAP2 (Figure 2G, H, S1). αGLAST-MNDs therefore reach and bind the astrocyte-restricted GLAST epitope *in vivo* that the free antibody labels. Together, these results indicate that αGLAST-MNDs preferentially target astrocytes in the intact brain.

**Figure 2.**
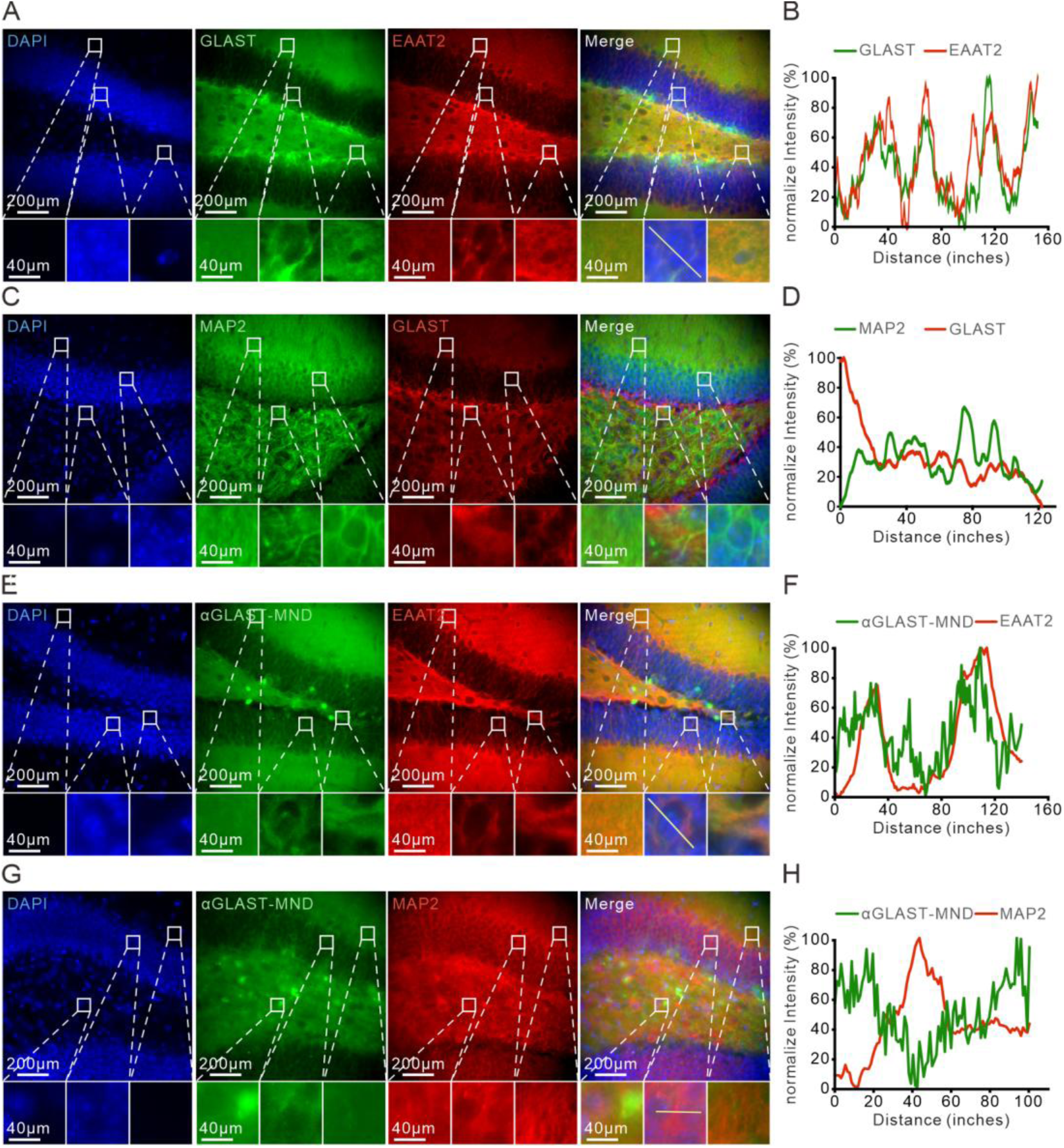
αGLAST-MNDs bind astrocyte-restricted GLAST epitopes in the dentate gyrus. (**A**) Fluorescence image of co-stained DAPI (blue), GLAST (green), and EAAT2 (red) in the DG. Bottom, enlarged images from the boxed regions. (**B**) Line-scan intensity profile of GLAST and EAAT2 along the line in the merged inset of (A). (**C**) Fluorescence image of co-stained DAPI (blue), MAP2 (green), and GLAST (red) in the DG. Bottom, enlarged images from the boxed regions. (**D**) Line-scan intensity profile of MAP2 (green) and GLAST (red) along the line in the merged inset of (C). (**E**) Fluorescence image of co-stained DAPI (blue), αGLAST-MND (green), and EAAT2 (red) in the DG. Bottom, enlarged images from the boxed regions. (**F**) Line-scan intensity profile of αGLAST-MND (green) and EAAT2 (red) along the line in the merged inset of (E). (**G**) Fluorescence image of co-stained DAPI (blue), αGLAST-MND (green), and MAP2 (red) in the DG. Bottom, enlarged images from the boxed regions. (**H**) Line-scan intensity profile of αGLAST-MND (green) and MAP2 (red) along the line in the merged inset of (G).

### 3.3 Low-amplitude AMFs evoke cell-type-specific astrocytic Ca²⁺ responses

To test whether magnetomechanical actuation of membrane-tethered MNDs drives astrocyte responses in the intact brain, cell-type-restricted GCaMP6 expression was combined with chronic fiber-photometry at hippocampal DG (Figure 3A). Astrocyte Ca²⁺ was recorded from mice expressing AAV5-gfaABC1D-GCaMP6f (astrocyte-restricted; viral delivery > 3 weeks pre-injection; Figure S2); neuronal Ca²⁺ was recorded from Thy1-GCaMP6s transgenic mice. Following stereotactic injection of αGLAST-MND or αGLAST-HND particles into the DG and implantation of the optical fiber above the injection site, fiber-photometry sessions were conducted weekly. During each session, AMF at 10 Hz and 25, 28, 40 or 50 mT were delivered as randomised trials.

**Figure 3.**
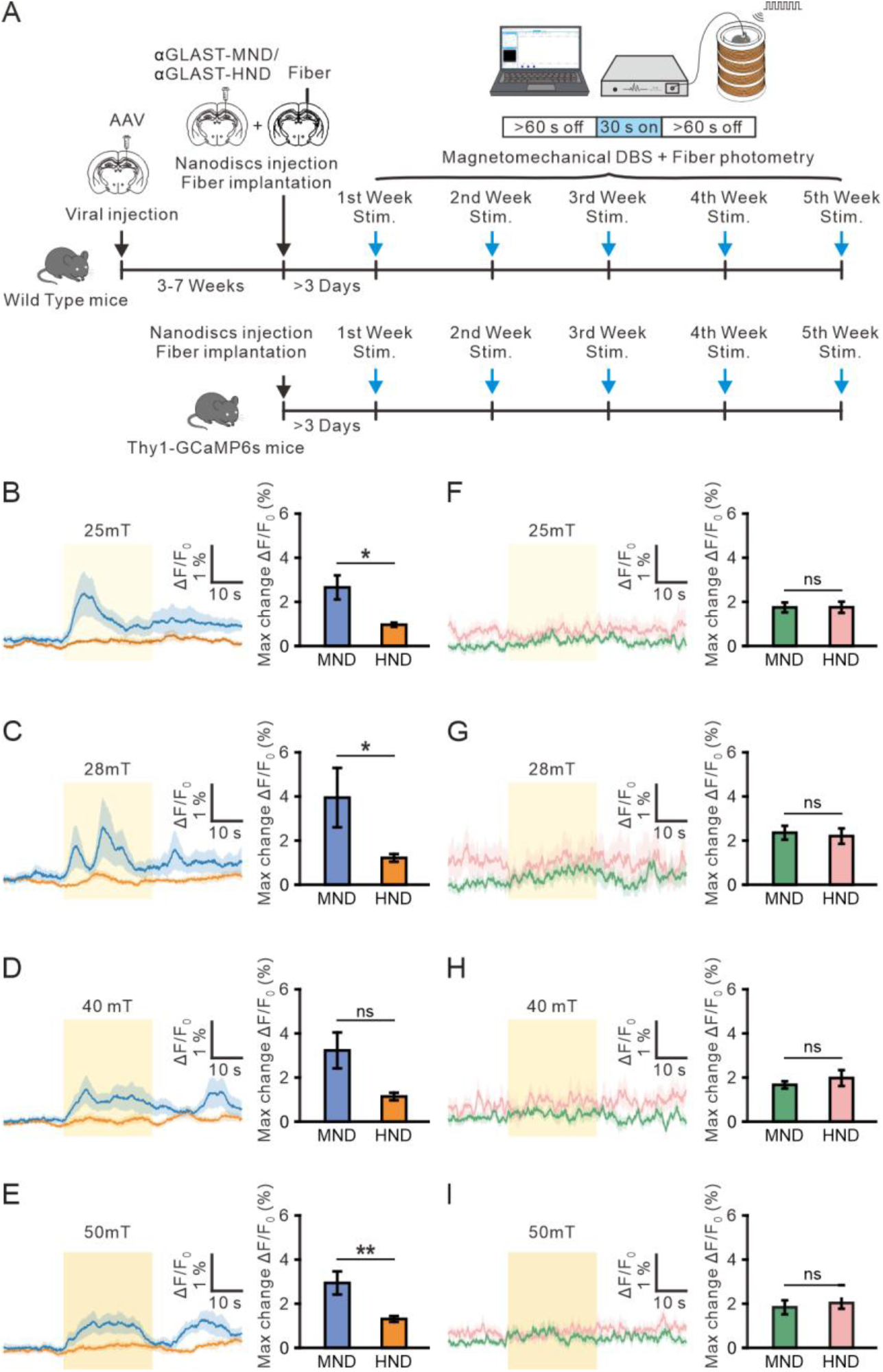
Cell-type-specific astrocytic Ca²⁺ responses evoked by αGLAST-MND stimulation across the 25 to 50 mT dose range. (**A**) Schematic of the αGLAST-MND stimulation and fiber-photometry paradigm. (**B** to **E**) Astrocyte cohort (gfaABC1D-GCaMP6f) at 25 mT (B), 28 mT (C), 40 mT (D) and 50 mT (E). Left, averaged ΔF/F₀ trace (blue, αGLAST-MND; orange, αGLAST-HND; light blue and light orange, SEM); yellow shading, 30 s AMF epoch. Right, max change ΔF/F₀. (**F** to **I**) Neuron cohort (Thy1-GCaMP6s) at 25 mT (F), 28 mT (G), 40 mT (H) and 50 mT (I). Left, averaged ΔF/F₀ trace (green, αGLAST-MND; pink, αGLAST-HND; light green and light pink, SEM); yellow shading, 30 s AMF epoch. Right, max change ΔF/F₀. Traces and per-mouse aggregation pooled across 2nd to 5th weeks post-injection. *p < 0.05; **p < 0.01; ns, no significance; Mann-Whitney U test.

To avoid potential influence of abnormal responses by surgery-induced reactive astrocytes during the 1st week (days 4-7 post-injection) [35, 36], AMF-evoked astrocyte Ca²⁺ responses were pooled across 2nd to 5th weeks post-injection, when the recording cohort was fully instrumented and had recovered from surgery (Figure 3). Representative averaged ΔF/F₀ traces show that αGLAST-MND stimulation produced reproducible Ca²⁺ responses (Figure 3B–E). In contrast, only weak responses were observed with αGLAST-HND control (Figure 3B–E). Quantification of peak-response amplitude revealed significantly larger αGLAST-MND-than αGLAST-HND-evoked responses at 25, 28 and 50 mT (Cohen d = +1.03 to +1.26, all p ≤ 0.022); a large effect at 40 mT did not reach statistical significance (d = +1.04, p = 0.109) (Figure 3B–E). Across the four field amplitudes, αGLAST-MND produced 2.2-to 3.3-fold larger peak responses than HND (MND +2.94 to +3.95% vs HND +1.14 to +1.31% ΔF/F₀). Cell-type specificity was tested by applying the identical stimulation protocol to Thy1-GCaMP6s mice with nanomaterials injection, in which the reporter labels neurons but not astrocytes. In the Thy1-GCaMP6s mice, peak Ca²⁺ responses did not differ significantly between αGLAST-MND and αGLAST-HND at any of the four field amplitudes when pooled across 2nd to 5th weeks (per-mT MND +1.44 to +1.84% vs HND +1.52 to +1.83%; Cohen d ranged −0.45 to +0.19; all p ≥ 0.59; Figure 3F–I). The MND-vs-HND effect was significantly larger in astrocytes (+2.15% ΔF/F₀) than in neurons (−0.08% ΔF/F₀; two-way ANOVA Material × Cell-type interaction F(1,25) = 4.44, p = 0.045; Cohen d = +1.60 [95% CI +0.88, +2.72]), demonstrating cell-type-specific astrocyte activation.

### 3.4 Stimulation efficacy is sustained for at least 5 weeks from a single injection

To analyze whether the magnetomechanically-induced glial response is stable over weeks, Ca2+ activity was quantified at each post-injection week for both cohorts (Figure 4, S3). The 1st week (days 4-7 post-injection) showed no clear material-specific effect in either cohort at any field amplitude (Figure 4A, F-I, S3; per-week Cohen d = −0.12 to +0.48), consistent with residual post-surgical recovery. From the 2nd week onward, the αGLAST-MND-mediated Ca²⁺ response emerged and remained consistently larger than that of the αGLAST-HND control at every field amplitude (Figure 4B-I, S3; per-week Cohen d = +0.70 to +2.83, median approximately +0.9), with no significant temporal trend across the observation window (linear regression of per-week Cohen d vs week for the 2nd to 5th weeks: slope = +0.19 per week, p = 0.10). Of the 16 (week × mT) combinations analyzed between the 2nd and 5th weeks, 14 reached bootstrap CI-based significance [30] and an additional 1 showed large effect sizes (|d| ≥ 0.7), yielding 15 of 16 combinations with large-effect MND advantage. This stable operating plateau extended to the 6th and 7th weeks in a smaller follow-up cohort (Figure S4).

**Figure 4.**
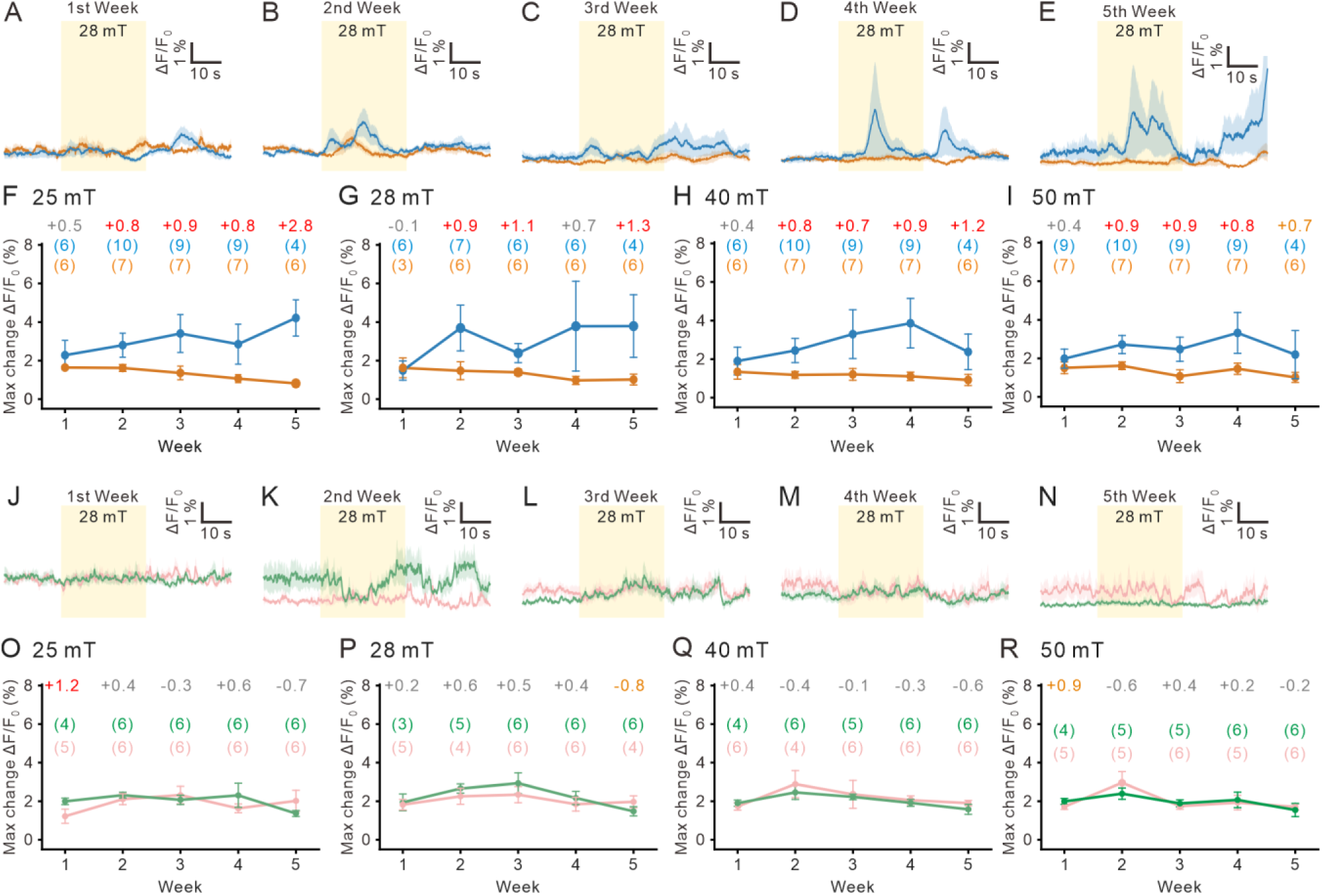
Cell-type-specific astrocyte response across 1st to 5th weeks post-injection. (**A** to **E**) Astrocyte cohort (gfaABC1D-GCaMP6f) averaged ΔF/F₀ traces at 28 mT for 1st week (A), 2nd week (B), 3rd week (C), 4th week (D) and 5th week (E), MND (blue) vs HND (orange). Mean ± SEM; yellow shading, 30 s AMF epoch. (**F** to **I**) Astrocyte per-week max change ΔF/F₀ (%) at 25 mT (F), 28 mT (G), 40 mT (H) and 50 mT (I), MND (blue) vs HND (orange). (**J** to **N**) Neuron cohort (Thy1-GCaMP6s) averaged traces at 28 mT for 1st week (J) to 5th week (N), MND (green) vs HND (pink). (**O** to **R**) Neuron per-week max change ΔF/F₀ (%) at 25 mT (O), 28 mT (P), 40 mT (Q) and 50 mT (R); layout as in F–I, MND (green) vs HND (pink). Cohen d annotated above each week (red, 95 % CI excludes 0; orange, |d| ≥ 0.7 but CI includes 0; grey, |d| < 0.7). Number of mice annotated above each week (blue or green, αGLAST-MND groups; orange or pink, αGLAST-HND groups).

The neuron cohort with Thy1-GCaMP6s mice (Figure 4J–R, S3D-F) showed no material-specific effect across any of the 5 weeks or four field amplitudes (per-week Cohen d distributed symmetrically around zero, median approximately +0.2, range −0.8 to +1.2; only 1 of 20 week × mT combinations reaching CI-based significance), confirming the absence of a neuronal response to the αGLAST-MND platform. Together, the per-week dataset establishes that a single intracranial injection of αGLAST-MND supports at least 5 weeks of stable, cell-type-specific astrocyte activation, extending to 7 weeks in a smaller follow-up cohort (Figure S4).

### 3.5 Local c-Fos induction is absent at low-amplitude stimulation

To examine whether the 28 mT × 10 Hz αGLAST-MND stimulation protocol drives transcriptional markers of activity at the injection site, c-Fos immunohistochemistry was performed on DG sections collected 90 min after a 10-cycle, 30 s on / 30 s off stimulation session in freely-moving mice (Figure 5A). The 28 mT amplitude was selected because it produced the strongest pooled MND-mediated Ca²⁺ response (Figure 3) and matched the field previously used for magnetomechanical stimulation of the subthalamic nucleus (STN) in a Parkinsonian mouse model [23]. The 3rd week post-injection (days 15-21) was chosen because it is safely past the reactive astrogliosis window that peaks between days 4 and 14 post-injection (1st and 2nd weeks) [35, 36]. To disentangle cell-type-specific from material-driven effects, four particle conditions were injected unilaterally into the DG: αGLAST-MND, αGLAST-HND, αThy1-MND and αThy1-HND; the contralateral DG served as the internal per-animal reference. Representative images confirmed the absence of stimulation-locked ipsilateral c-Fos induction for αGLAST (Figure 5B-D) and αThy1 (Figure 5E-G) conditions. The ipsilateral / contralateral c-Fos ratio did not differ significantly between MND and HND for either the anti-GLAST (t = +1.68, p = 0.12; Figure 5D) or the anti-Thy1 targeting antibody (t = −1.63, p = 0.13; Figure 5G). αGLAST-MND was the only condition whose ratio approached unity (1.00), whereas the other three conditions ranged from 0.80 to 0.87. The null c-Fos result is the expected outcome given that αGLAST-MND stimulation does not evoke a neuronal Ca²⁺ response (Figures 3F– I, 4J–N and 4O–R), and independently supports the cell-type-specific engagement of the platform.

**Figure 5.**
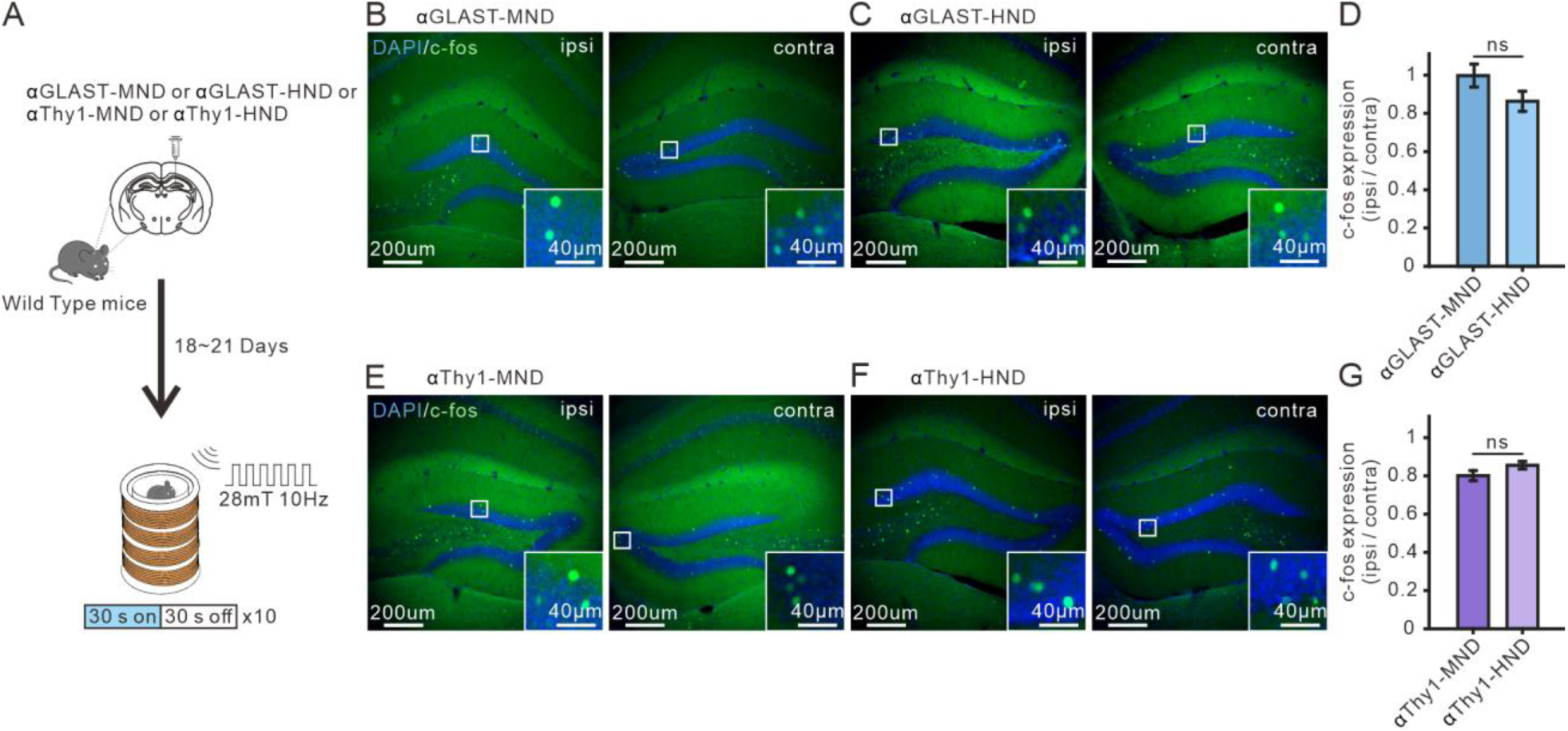
Low-amplitude AMF stimulation did not induce c-Fos expression at the injection site for any of the four particle conditions. (**A**) Schematic of the c-Fos experiment. Wild-type mice received a unilateral stereotactic injection of one of four particle conditions into the right dentate gyrus (αGLAST-MND, αGLAST-HND, αThy1-MND or αThy1-HND). At 3rd week, mice were stimulated with a 10 Hz AMF at 28 mT delivered as 10 cycles of 30 s on / 30 s off, then perfused for c-Fos immunohistochemistry. (**B** and **C**) Representative c-Fos expression (green, c-Fos; blue, DAPI) of DG from αGLAST-MND (B) and αGLAST-HND (C) conditions, comparing ipsilateral (left) and contralateral (right) hemispheres of the same animal. Insets, high-magnification views. (**D**) c-Fos-positive nuclear density ratio (ipsi/contra) per animal for the αGLAST-conjugated nanomaterials conditions (n = 6 mice per group). Independent-samples t-test MND vs HND: t = +1.68, p = 0.12; ns, no significance. (**E** and **F**) Representative c-Fos expression (green, c-Fos; blue, DAPI) of DG from αThy1-MND (E) and αThy1-HND (F) conditions, comparing ipsilateral (left) and contralateral (right) hemispheres of the same animal. Insets, high-magnification views. (**G**) c-Fos-positive nuclear density ratio (ipsi/contra) per animal for the αThy1-conjugated nanomaterials conditions (n = 6 mice per group). Independent-samples t-test MND vs HND: t = −1.63, p = 0.13; ns, no significance.

## 4. Discussion

We demonstrate that antibody-functionalized magnetite vortex nanodiscs (αGLAST-MND) enable wireless, transgene-free, cell-type-specific stimulation of native hippocampal astrocytes. Under 10 Hz AMF stimulation, αGLAST-MND evoked astrocyte Ca²⁺ responses 2-to 3-fold larger than non-magnetic controls and 2.1-fold larger than paired neuronal responses from Thy1-GCaMP6s mice, with no material-specific response in the neuron cohort (Figure 3). A single intracranial injection sustained stable astrocyte responses across the 5-week observation window and extended to 7 weeks in a smaller cohort (Figure 4, S4). The low field amplitude (<30 mT) and affordable, scalable magnetic system [25] make the platform readily suitable for freely behaving experiments and open opportunities for broader clinical and translational applications.

The αGLAST-MND platform builds on the prior anti-GLAST-targeted magnetomechanical astrocyte stimulation study [24], which established the feasibility of engaging endogenous astrocyte mechanosensors through antibody-targeted iron oxide particles. The critical distinction between the two approaches lies in both nanoparticle physics and stimulation hardware. The iron oxide spheres used previously displaced membrane-bound particles via magnetic gradient force, which requires a steep spatial gradient in the applied field [24]. Achieving sufficient gradient therefore required either the fringe field of a 9.4 T MRI scanner or a purpose-built permanent magnet array, restricting deployment to institutions with dedicated imaging or magnet infrastructure. Magnetite vortex nanodiscs instead generate torque in a uniform alternating magnetic field, without requiring any spatial field gradient [20, 22, 31]. Their flux-closed vortex ground state combined with high saturation magnetization (Ms ≈ 135 Am² kg⁻¹) and a low coercive field (∼27 mT) produces substantial torque per particle at low field amplitudes [20, 22, 31]. Prior calculations for a 226-nm vortex disc at 26 mT estimated ∼18 pN of force per particle at the disc edge, well above the ∼0.3 pN activation threshold of mechanosensors such as TRPV4 and Piezo1 [20]. This regime is achievable at the 25 to 28 mT deliverable by a bench-top open-source coil system [25]. The low-field competence of the vortex particle is what enables the affordable, scalable coil hardware. The coil design can be built from commodity components, replicated across laboratories, and reconfigured for different chamber sizes and freely-behaving recording setups, supporting routine multi-laboratory use and future translational adaptation.

A recent study reported behavioral recovery in the Parkinsonian mouse model using non-antibody-conjugated MNDs at a comparable particle dose in the STN and stimulation at 28 mT, 5 Hz [23]. That study reported c-Fos induction in downstream motor targets but did not report c-Fos at the STN injection site. Another study shows that at the higher amplitude of 50 mT at 10 Hz, non-antibody-conjugated MND stimulation activates neuronal TRPC channels and induces local c-Fos in STN neurons [22], indicating that neuronal recruitment is amplitude-dependent. Two factors likely account for the absent αThy1-MND neuronal c-Fos observed at 28 mT × 10 Hz in the dentate gyrus. First, hippocampal granule cells may differ from STN neurons in mechanosensor expression or coupling. Second, low-amplitude stimulation at 28 mT is a regime in which local IEG induction at the injection site is difficult to resolve [23]. The convergent 28 mT field regime engages endogenous mechanosensitive channels across multiple brain regions and cell types. Whether astrocyte and neuron mechanosensor recruitment can be selectively balanced through field parameter tuning remains to be determined. Sustained αGLAST-MND-mediated Ca²⁺ response for 5 to 7 weeks from a single intracranial injection represents a substantial advance from acute proof-of-concept.

Hippocampal astrocytes in freely behaving mice exhibit continuous spontaneous Ca²⁺ activities [37, 38]. Combined with our baseline-selection procedure (AMF onset was manually triggered only after real-time inspection of the photometry trace confirmed a stable, quiet pre-stimulus baseline), this ongoing activity predicts that a proportion of trials will contain a spontaneous transient in the 30 s post-onset window and register as an apparent increase above the intentionally quiet baseline, regardless of the material injected. Consistent with previous research, we found a small Ca²⁺ response in the αGLAST-HND group (max change ΔF/F₀ ∼1.2%). The αGLAST-HND cohort therefore serves as a matched-condition reference for baseline-selection-related spontaneous activity, and the systematic 2.2-to 3.3-fold larger αGLAST-MND-mediated Ca²⁺ response above this reference reflects the MND-driven component of the signal.

In addition, intracranial injection can transiently induce reactive astrogliosis, which is characterized by elevated resting Ca²⁺ activity and more frequent spontaneous transients in vivo [39, 35]. We found that the 1st week (days 4-7 post-injection) profile differs qualitatively from later time points (Figure 4F–I, S3). In the 1st week, the spontaneous response in the αGLAST-HND group was relatively large (max change ΔF/F₀ 1.35 to 1.64% across the four amplitudes) and the two materials were statistically indistinguishable (per-week Cohen d = −0.12 to +0.48; Mann-Whitney p ≥ 0.71 at all field intensities). By the 5th week, the Ca²⁺ response in the αGLAST-HND group had declined by 31 to 50% (to 0.82 to 1.02%) while the αGLAST-MND-mediated Ca²⁺ response rose to 2.20 to 4.22%; the between-material effect increased to Cohen d = +0.73 to +2.83 and reached statistical separation at 25 and 28 mT (Mann-Whitney p ≤ 0.038; Figure 4F–I, S3). This trajectory reflects the sub-acute reactive phase captured during the 1st week (days 4-7 post-injection), which enlarges the αGLAST-HND baseline and masks the material-specific response, and its subsequent resolution, which restores a quieter baseline that reveals the αGLAST-MND-mediated Ca²⁺ response.

Although the combined fiber-photometry and c-Fos data support preferential astrocyte activation without a statistically significant neuronal response, we cannot fully exclude the possibility of low-level off-target MND binding to non-astrocytic epitopes below the detection thresholds of the assays used here. Second, the downstream molecular mechanism of MND-mediated stimulation remains unresolved; TRPV4, TRPC1/5/6 and Piezo1 are all plausible candidates implicated across prior magnetomechanical astrocyte studies [24, 26, 27, 28, 29], and pharmacological dissection is necessary to clarify the underlying mechanism. Third, anti-GLAST was chosen for its established astrocyte specificity, but potential functional side effects on glutamate transport or GLAST-associated signaling have not been systematically characterized; translational deployment will require detailed safety assessment of anti-GLAST antibody or the identification of alternative astrocyte-specific membrane targets that avoid perturbing endogenous function. Finally, while the platform’s wireless, cell-type-specific stimulation capability is established, behavioral and therapeutic outcomes in disease models remain to be demonstrated.

Beyond the current demonstrations, wireless AMF stimulation is compatible with freely-behaving assays that are constrained by tethered optical fibers, such as social behavior, complex motor coordination, and long-duration observation. The antibody-based targeting layer is potentially extensible: substituting anti-GLAST with monoclonal antibodies against other astrocyte-selective epitopes could tune the platform to specific astrocyte subtypes, and the same particle chemistry could be redirected to microglia, oligodendrocytes, or defined neuronal populations by antibody swap. The most immediate translational tests are in chronic disease models where astrocyte activation itself is therapeutic, including Parkinsonian motor recovery [13], memory rescue in Alzheimer’s disease models [2], and pain modulation [7]. The scalable magnetic system [25] can be accommodated to different animal sizes without redesigning the particle chemistry, making extension to rat and larger animal models tractable.

### Conclusion

We have engineered and validated a wireless, transgene-free, cell-type-specific DBS platform based on antibody-targeted magnetite vortex nanodiscs. Magnetomechanical actuation by low-amplitude AMF (25 to 28 mT) combined with anti-GLAST antibody targeting activates native astrocytes through endogenous mechanosensors without genetic modification or chronically implanted electronic hardware. Stimulation was delivered by an open-source bench-top coil system [25]. This hardware supports deployment in awake, freely behaving animals without MRI infrastructure or permanent magnet arrays. A single intracranial injection sustained astrocyte responses for at least 5 weeks and up to 7 weeks in a smaller cohort. The platform offers a wireless magnetomechanical route to astrocyte-targeted neuromodulation for basic research, with potential clinical applications where enhancing astrocyte activity may carry therapeutic benefits.

## Author contributions

**Yi-Ting Cai:** Data Curation, Methodology, Investigation, Formal analysis, Visualization, Validation, Writing – review & editing. **Jiao-Cheng Wang**: Data Curation, Formal analysis. **Po-Han Chiang**: Supervision, Conceptualization, Methodology, Software, Formal analysis, Investigation, Visualization, Funding acquisition, Project administration, Writing – Original draft, Writing – Review & Editing.

## Declaration of competing interest

The authors declare that they have no known competing financial interests or personal relationships that could have appeared to influence the work reported in this paper.

## Acknowledgements

We thank Prof. Tsai-Wen Chen (National Yang Ming Chiao Tung University) for kindly providing the Thy1-GCaMP6s transgenic mice. We are also grateful to Prof. Albert C. Yang (National Yang Ming Chiao Tung University) for insightful discussion of the study and for assistance with funding acquisition.

## Funding

We thank National Science and Technology Council, Taiwan for Funding this study. NSTC 113-2321-B-A49-020; NSTC 114-2321-B-A49-003; NSTC 114-2321-B-A49-014; NSTC 113-2320-B-A49-044; NSTC 114-2320-B-A49-027; NSTC 114-2634-F-A49-006

## Data availability statement

The data that support the findings of this study are available from the corresponding author upon reasonable request.

## Ethics statement

All animal procedures were approved by the Institutional Animal Care and Use Committee (IACUC) of National Yang Ming Chiao Tung University under protocol number IACUC NO. 114009 (PI: Prof. Po-Han Chiang), and were conducted in accordance with the Guide for the Care and Use of Laboratory Animals of NYCU. The experimental design, procedures, and reporting adhered to the ARRIVE 2.0 guidelines [40]. Consistent with the 3Rs principles of Replacement, Reduction, and Refinement, sample sizes were minimised while retaining sufficient statistical power, and postoperative analgesia and daily monitoring were provided to reduce animal suffering. Full details of housing conditions, surgical procedures, and euthanasia are provided in the Materials and Methods and Supplementary Methods.

## Declaration of generative AI and AI-assisted technologies in the manuscript preparation process

During the preparation of this work, the author used Claude (Anthropic), Gemini, and ChatGPT for assistance with manuscript editing and code development for data analysis. The authors reviewed and verified all outputs and took full responsibility for the content of this work.

## Notes

### Competing Interest Statement

The authors have declared no competing interest.

