## Supplementary Methods for "Astrocyte-Selective Wireless Deep Brain Stimulation by Magnetite Nanodiscs at Low Alternating Magnetic Field"

### Content:

Supplementary Methods

Supplementary Figures: S1 to S4.

### Supplementary Methods

#### S1 Synthesis of hematite and magnetite nanodiscs

Hematite nanodiscs were synthesised by combining 0.8 g sodium acetate anhydrous, 10 mL ethanol (99 %), 600 μL dd$H_2O$ and 0.273 g iron(III) chloride hexahydrate ($FeCl_3 \cdot 6H_2O$) in a Teflon-lined reaction vessel, which was sealed in a stainless-steel autoclave and heated at 180 °C for 18 h. After cooling to room temperature, the product was collected by centrifugation (8,500 rpm, 5 min), washed three times with dd$H_2O$ and three times with ethanol (99 %), dried under vacuum and stored in hexane. Magnetite nanodiscs were prepared by reduction of the hematite nanodiscs. Briefly, 100 mg of hematite nanodiscs were mixed with 20 mL trioctylamine (97 %) and 1 g oleic acid in a reactor connected to a gas supply system. The reactor was purged with nitrogen (99.9 %) followed by a 5 % $H_2$ / 95 % Ar mixture and heated at 370 °C for 25 min. The reduced product was collected by centrifugation (8,500 rpm, 5 min), washed three times with hexane, dried under vacuum and stored in chloroform until use.

#### S2 PMAO coating and antibody conjugation

For cell-type-specific stimulation, nanodiscs were surface-modified with poly(maleic anhydride-alt-1-octadecene) (PMAO) and subsequently conjugated with anti-GLAST or, as a targeting control, anti-Thy1 monoclonal antibody. One mg of nanodiscs and 10 mg of PMAO were combined in 1 mL chloroform and sonicated for 1 h, then dried under vacuum. The dried PMAO-coated nanodiscs (5 mg) were resuspended in 8 mL of 10× TAE buffer and incubated at 80 °C with continuous sonication for 3 h, then collected by centrifugation (8,500 rpm, 5 min) and washed three times with dd$H_2O$. For antibody conjugation, the PMAO-coated nanodiscs were resuspended in 0.1 M MES buffer with 0.5 M NaCl; surface carboxyl groups were activated by EDC and NHS at final concentrations of 2 mM and 5 mM, respectively, for 15 min at room temperature. The activated nanodiscs were washed three times with PBS, resuspended in PBS, and incubated with anti-GLAST antibody (rat monoclonal, clone ACSA-1; Miltenyi Biotec #130-095-822; 1 μL per mg of nanodisc) overnight at 4 °C. Antibody-conjugated nanodiscs were collected by centrifugation, washed three times with PBS to remove unbound antibody, and resuspended in PBS at 2 mg $mL^{-1}$ for subsequent experiments. Hematite nanodiscs (HNDs) bearing the same antibody coating but lacking magnetic remanence were prepared in parallel and served as the magnetic-effect control throughout all in vivo experiments. anti-Thy1 antibody (clone G7, catalog #14-0901-82) was used at the same 1 μL per mg of nanodisc ratio for c-Fos control experiments.

#### S3 Material characterization

Particle morphology and dimensions were assessed by transmission electron microscopy (TEM; HT7800, Hitachi) at the Microscopy Core Laboratory of Chang Gung Memorial Hospital, operated at 100 kV, on samples drop-cast onto carbon-coated copper grids (Ted Pella Inc.). Particle diameter was quantified from ≥ 6 particles per batch using ImageJ. Magnetic properties were characterized by vibrating-sample magnetometry (MPMS SQUID, Quantum Design) at the Core Facility Center of National Cheng Kung University at room temperature with a maximum applied field of ± 0.9 T. Crystal phase was identified by powder X-ray diffraction (XRD; XtaLAB Synergy DW, Rigaku, equipped with a HyPix-Arc 150° curved hybrid photon-counting detector and a graphite monochromator) at the Instrumentation Center of National Tsing Hua

University. Hydrodynamic diameter, polydispersity and zeta potential of PMAO-coated and antibody-conjugated nanodiscs were measured by dynamic and electrophoretic light scattering (DelsaNano C Particle Analyzer, Beckman Coulter) in $ddH_2O$. Iron content of nanodisc suspensions was quantified by inductively coupled plasma optical emission spectrometry (ICP-OES; Agilent 725) at the National Cheng Kung University Core Facility Center.

**S4 Animal experiments**

Adult male C57BL/6J mice (BioLASCO Taiwan) were 4 to 7 months old at the time of viral injection for the fiber-photometry cohort and 4 to 5 months old for the c-Fos cohort. Adult male Thy1-GCaMP6s transgenic mice (Jackson Laboratory, strain #024275) were 7 to 14 months old at the time of surgery. Mice were group-housed under a 12 h light / dark cycle in a temperature- and humidity-controlled vivarium with food and water available ad libitum. All experimental procedures were approved by the National Yang Ming Chiao Tung University (NYCU) Institutional Animal Care and Use Committee (IACUC protocol #114009) and reported in accordance with the ARRIVE 2.0 guidelines. Mice were randomly assigned to MND or HND treatment groups using a computer-generated random sequence. Experimenters performing surgery and photometry were not blinded to group identity, but signal processing and IHC quantification were performed by analysts blinded to group assignment.

**S5 Stereotactic surgery and viral injection**

Mice were anaesthetised with isoflurane (3.5 % for induction, 1 to 2 % v / v in 1 L $min^{-1}$ $O_2$ for maintenance) and secured in a stereotactic frame. Body temperature was maintained at 37 °C with a feedback-controlled heating pad and ophthalmic ointment was applied. Coordinates were determined relative to bregma and lambda. For astrocyte-restricted $Ca^{2+}$ imaging, AAV5-pZac2.1-gfaABC1D-cyto-GCaMP6f ($\geq 7 \times 10^{12}$ vg $mL^{-1}$; Addgene #52925 or Vector Biolabs custom prep) was pressure-injected (0.6 µL per site) into the dentate gyrus (AP −1.9 mm, ML +1.2 mm, DV −1.9 mm) at 100 nL $min^{-1}$ using a microsyringe. After at least 3 weeks of viral expression, αGLAST-MND or αGLAST-HND particles (2 mg $mL^{-1}$, 2 µL) were injected at the same coordinates. An optical fiber (200 µm core diameter, 3 mm length, 0.48 NA) was immediately implanted 100 µm dorsal to the injection site (DV −1.8 mm) and secured to the skull with dental adhesive resin (Super-Bond, Sun Medical). Postoperative analgesia (carprofen, 5 mg $kg^{-1}$ subcutaneous) was administered for three consecutive days. The first magnetic stimulation session was conducted no earlier than 3 days after implantation.

**S6 Magnetic stimulation apparatus**

The magnetic stimulation system followed the open-source air-core coil design previously described [25]. Briefly, four air-core coils were wound with approximately 350 turns of 12-AWG enamelled copper wire on a cylindrical former with an inner bore that accommodated the mouse stimulation chamber. The measured resistance and inductance of individual coils were 1.02 to 1.55 Ω and 22 to 31.6 mH, respectively. AMFs were generated by full-bridge driver modules (AQMH3615NS, AKELC) driven by 5 V square-wave signals from an Arduino UNO microcontroller (Arduino) programmed via the Arduino IDE with custom scripts to deliver 10 Hz field oscillations at nominal amplitudes of 25, 28, 40 or 50 mT. Coils were powered by an external DC power supply (HJS-1000, Huntkey), and the peak magnetic-field amplitude at the animal position was calibrated before every experimental cohort with a calibrated Hall-effect gaussmeter (TM801, KANETEC).

**S7 Fiber-photometry recording**

Fiber-photometry was performed using a one-fiber-photometry system in freely moving, awake mice within the magnetic stimulation chamber. GCaMP fluorescence was excited alternately at 410 nm (isosbestic control, 38.11 % LED intensity) and 470 nm ($Ca^{2+}$-dependent signal, 17.18 % LED intensity); emission was collected through a 525 / 30 nm bandpass filter, with 410 nm and 470 nm channels interleaved at a total acquisition rate of 30 frames per second (15 fps per channel). On each recording day, mice were habituated to the experimental room for 1 h and to the stimulation chamber for a further 10 min before signal acquisition. A ≥ 2 min baseline recording was obtained at the start of every session. The alternating magnetic field was manually triggered by the experimenter, based on real-time inspection of the ongoing photometry trace, only when the $\Delta F/F_0$ signal was stable and the animal was relatively stationary for at least 20 s prior to onset. This procedure ensured a quiet pre-stimulus baseline for every trial. Each stimulation trial delivered 30 s of AMF (10 Hz at 25, 28, 40 or 50 mT) followed by a 60 s post-stimulation recording window and a rest interval of at least 60 s before the next trial, with up to 8 randomised trials per session. Fiber-photometry recordings were performed once or twice per week, starting from day 4 post-injection. To exclude the acute post-surgical window (days 1-3), recording sessions were grouped into weekly windows relative to injection: 1st week (days 4-7), 2nd week (days 8-14), 3rd week (days 15-21), 4th week (days 22-28), 5th week (days 29-35), 6th week (days 36-42) and 7th week (days 43-49).

**S8 Photometry signal processing**

Raw 470 nm fluorescence traces were exported and processed in Python (v 3.9.0) using pandas, numpy and scipy. A 5 Hz low-pass Butterworth filter was applied to attenuate high-frequency shot noise. For each trial the recording was aligned to the AMF onset (t = 0 s) and a per-trial normalized trace was computed as $\Delta F/F_0 = (F - F_0) / F_0$, where $F_0$ is the mean of the lowest 5% of samples within the preceding 100 s window. Two analysis time windows were defined for each trial: a 20 s baseline window (−20 to 0 s relative to AMF onset) and a 30 s stimulation window (0 to +30 s). The primary metric was the max change $\Delta F/F_0$, defined as the maximum $\Delta F/F_0$ during the 30 s stimulation window minus the mean $\Delta F/F_0$ across the preceding 20 s baseline window. All $\Delta F/F_0$ values throughout the manuscript and figure captions are expressed as %. Trials in which the standard deviation of $\Delta F/F_0$ across the 20 s baseline window exceeded 0.65% were excluded from downstream analysis (250 of 1,969 traces, 12.7%; no mouse excluded). Trials with pronounced locomotion artifact on the 410 nm isosbestic channel were flagged and reviewed manually before inclusion.

**S9 Magnetic stimulation for c-Fos induction**

For the c-Fos experiment, four particle conditions were prepared: αGLAST-MND, αGLAST-HND, αThy1-MND and αThy1-HND (2 mg $mL^{-1}$, 2 µL each). Each condition was stereotactically injected unilaterally into the right dentate gyrus (AP −1.9 mm, ML +1.2 mm, DV −1.9 mm) of adult male C57BL/6J mice (n = 6 mice per group), with the non-injected contralateral hemisphere serving as the internal per-animal reference. Three weeks after injection, mice were acclimatised to the experimental room for 1 h, transferred to the cylindrical stimulation chamber for 10 min, and stimulated with an AMF at 28 mT, 10 Hz, delivered as 10 cycles of 30 s on / 30 s off. Ninety minutes after the end of the stimulation session, mice were transcardially perfused with PBS followed by 4 % paraformaldehyde and brain tissue was collected for immunohistochemistry.

**S10 Immunohistochemistry, image acquisition and quantification**

Brains were post-fixed in 4 % paraformaldehyde overnight at 4 °C, sectioned coronally at 50 µm on a vibratome (5100 MZ, Campden Instruments; amplitude 0.5, frequency 50 Hz), washed three times in PBS for 5 min each, and permeabilised in 2 % Triton X-100 / PBS for 15 min. To reduce endogenous fluorescence, sections were incubated in 10 % methanol, 3 % $H_2O_2$ and 2 % Triton X-100 in PBS for 10 min followed by three PBS washes. Sections were blocked with 3 % normal goat serum for 60 min at room temperature, washed three times in PBS, and incubated overnight (16-18 h) at 4 °C with the following primary antibodies diluted in PBS containing 1 % normal goat serum and 2 % Triton X-100: rabbit anti-c-Fos (1:500; clone 9F6, #2250, Cell Signaling Technology); rabbit anti-GFAP (1:750; clone E4L7M, #80788, Cell Signaling Technology); rabbit anti-EAAT2 (1:500; ab205248, Abcam); mouse anti-GLAST (1:750; #130-095-822, Miltenyi Biotec); chicken anti-MAP2 (1:750; ab5392, Abcam). After primary antibody incubation and three PBS washes, sections were incubated for 2 h at room temperature with Alexa Fluor-conjugated secondary antibodies (1:500 in PBS): rabbit Alexa 488 (ab150077, Abcam), rabbit Alexa 594 (ab150080, Abcam), mouse Alexa 488 (ab150113, Abcam), mouse Alexa 594 (ab150116, Abcam), chicken Alexa 488 (ab150173, Abcam), chicken Alexa 594 (ab150176, Abcam), or rat Alexa 594 (ab150160, Abcam). Sections were washed three times in PBS, mounted onto glass slides and coverslipped using Fluoroshield with DAPI (GeneTex). Images were acquired on an inverted fluorescence microscope (DMI3000, Leica) or an upright fluorescence microscope (SS-1000-00, Scientifica) equipped with an LED light source (pE-300, CoolLED), a Hamamatsu C13440 camera and filter cubes (39000, 19008 and 31002; Chroma Technology). c-Fos-positive nuclei were quantified within the dentate gyrus using automated thresholding in ImageJ (Fiji).

### Supplementary Figures

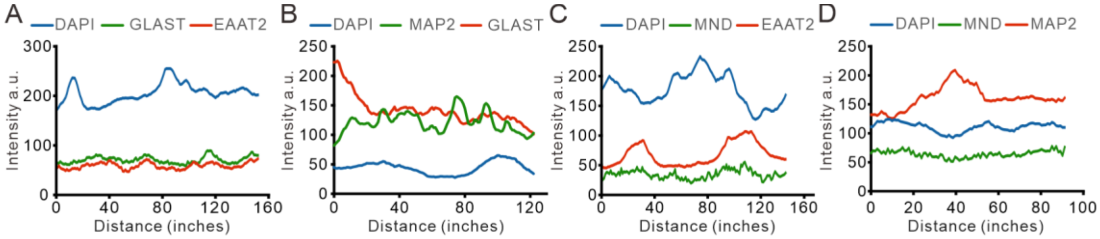


S1.6 Magnetic stimulation apparatus — coil specifications

**Supplementary Figure S1. Raw data of Figure2. Absolute fluorescence intensity along the four line-scans shown in Figure 2.** (**A**) DAPI (blue), GLAST (green) and EAAT2 (red) channels, corresponding to Figure 2A (astrocyte marker co-staining). (**B**) DAPI (blue), GLAST (green) and MAP2 (red) channels, corresponding to Figure 2C (astrocyte marker vs neuronal marker). (**C**) DAPI (blue), αGLAST-MND (green) and EAAT2 (red) channels, corresponding to Figure 2E. (**D**) DAPI (blue), αGLAST-MND (green) and MAP2 (red) channels, corresponding to Figure 2G (MND vs neuronal marker). Whereas Figure 2 shows per-channel-normalized intensities, panels here retain the raw arbitrary-unit intensities for reference.

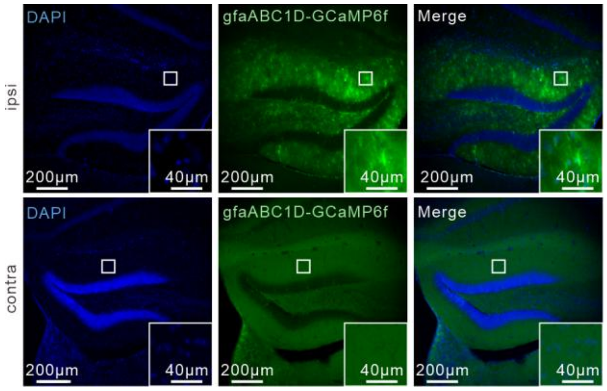


**Supplementary Figure S2. Expression of gfaABC1D-GCaMP6f.** Representative fluorescence images of gfaABC1D-GCaMP6f expression in the DG of a wild-type mouse three weeks after unilateral stereotactic injection of AAV5-gfaABC1D-GCaMP6f. Top row, ipsilateral hemisphere; bottom row, contralateral (non-injected) hemisphere. Left column, DAPI (blue); middle column, gfaABC1D-GCaMP6f native fluorescence (green); right column, merged. Insets, high-magnification views. Scale bars, 200 μm (main) and 40 μm (inset).

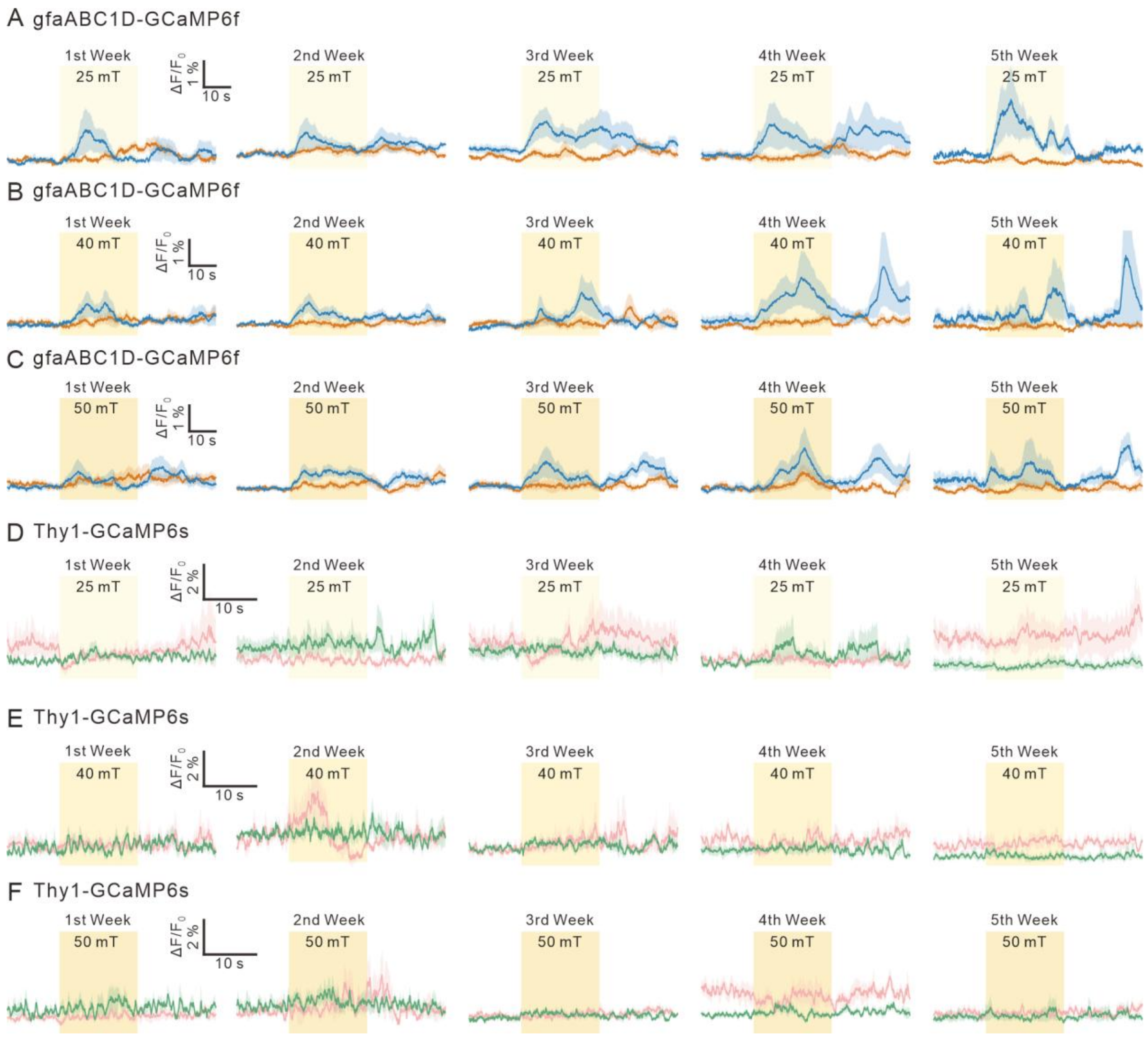


**Supplementary Figure S3. Per-week $\Delta F/F_0$ traces at 25, 40 and 50 mT (fields not shown in Figure 4).** Trial- and mouse-averaged $\Delta F/F_0$ traces organized by cohort and field amplitude (rows) and post-injection week (columns). (**A** to **F**) Astrocyte cohort (gfaABC1D-GCaMP6f): (A) 25 mT, (B) 40 mT and (C) 50 mT, MND (blue) vs HND (orange). Neuron cohort (Thy1-GCaMP6s): (D) 25 mT, (E) 40 mT and (F) 50 mT, MND (green) vs HND (pink). In each row, the five columns correspond to the 1st, 2nd, 3rd, 4th and 5th weeks post-injection. Light area, SEM; yellow shading, 30 s AMF epoch. n as reported in the corresponding per-week bar plots of Figure 4F–I (astrocyte) and 4O–R (neuron).

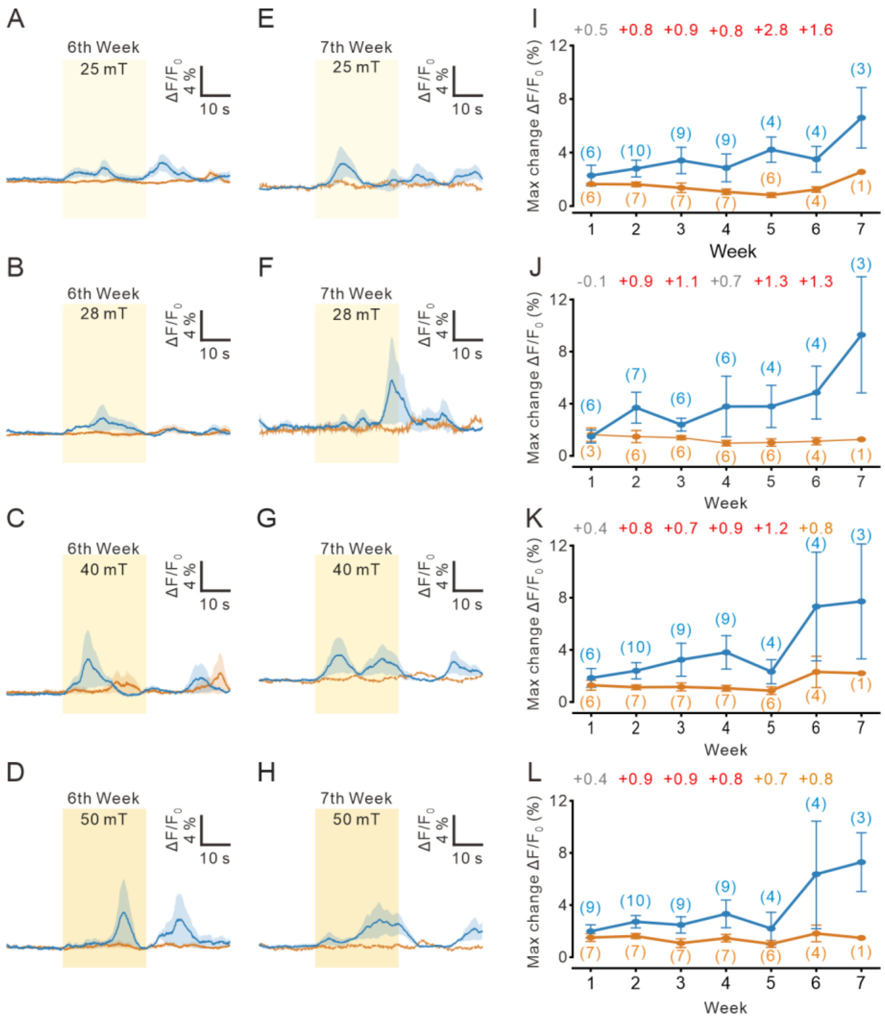


**Supplementary Figure S4. Extended astrocyte durability across 6th and 7th weeks post-injection.** (**A** to **D**) Trial- and mouse-averaged $\Delta F/F_0$ traces at the 6th week for 25 mT (A), 28 mT (B), 40 mT (C) and 50 mT (D), MND (blue) vs HND (orange). (**E** to **H**) Same as (A) to (D) but for the 7th week. Yellow shading, 30 s AMF epoch. (**I** to **L**) Per-week astrocyte max change $\Delta F/F_0$ (%) at 25 mT (I), 28 mT (J), 40 mT (K) and 50 mT (L), MND (blue) vs HND (orange), extended to the 7th week. Cohen d annotated above each week (red bold, 95 % CI excludes 0; orange bold, $|d| \geq 0.7$ but CI includes 0; grey, $|d| < 0.7$). Number of mice annotated at each week (blue, αGLAST-MND groups; orange, αGLAST-HND groups).
